# Accuracy of retrospective birth location data – An analysis based on siblings

**DOI:** 10.1101/2023.07.21.550064

**Authors:** Stephanie von Hinke, Nicolai Vitt

**Affiliations:** University of Bristol, Institute for Fiscal Studies; University of Bristol

## Abstract

Many surveys ask participants to retrospectively record their location of birth. This paper examines the accuracy of such data in the UK Biobank using a sample of siblings. Comparison of reported birth locations for siblings with different age gaps allows us to estimate the probabilities of household moves and of misreported birth locations. We find an annual probability of 1.2% for household moves of one kilometre or more, suggesting that geographical mobility during childhood was low. Our results furthermore show a sizeable probability of misreporting, with 28% of birth coordinates, 16% of local districts and 6% of counties of birth being incorrectly reported. We show that such error can lead to substantial attenuation bias when investigating the impacts of location-based exposures, especially when there is little spatial correlation and limited time variation in the exposure variable. Sibling fixed effect models are shown to be particularly vulnerable to the attenuation bias.

## Introduction

Retrospective data collection on residential location is common in secondary data sources. For example, many datasets include individuals’ residential location at birth or in early childhood, recollected by survey participants in adulthood or older age (e.g., the US *National Longitudinal Survey of Youth 1979*, the *UK Biobank* and *Understanding Society*, the German *Socio-Economic Panel*, the Dutch *Lifelines cohort*, and the Scottish *Generation Scotland*). These location data have in turn been used in a wide range of empirical applications, such as those studying geographic mobility^1,2^, geographic stratification and spatial correlation of genetic variation^3–8^, assortative mating and social homogamy^9,10^, but they have also been used to capture regional differences in infrastructure, health or economic circumstances, such as the staggered roll-out of policy^11^. Similarly, they have allowed researchers to include area of birth fixed effects to account for systematic differences between geographical areas^12,13^, and to merge in external information on (area-level) weather, health or socio-economic information^14–21^. Despite much research in a wide range of applications relying on these retrospectively recorded (birth) locations, very little work has explored the accuracy of these data, especially considering they often rely on individuals’ correct 30+ year recall. This is the aim of our paper.

We focus on the UK Biobank (UKB), a large cohort study of half a million individuals aged 45–69 between 2006–2010. It includes detailed environmental, lifestyle, health and genetic data, but has very limited information on the environment in which individuals grew up and the circumstances during their early childhood. It does, however, record individuals’ location of birth. These data are based on the following question which was asked by the interviewers to any participant who indicated being born in England, Scotland or Wales: *“What is the town or district you first lived in when you were born?”* Based on the respondent’s answer, the interviewer selected the corresponding place from a very long and detailed list of place names in the UK. The birth place was then converted into coordinates (eastings and northings at a 1 kilometre resolution) which are provided in the data.

This paper explores the accuracy of these data and highlights any potential implications of using such data for future research. We exploit the fact that the UKB includes a sample of approximately 40,000 full siblings which can be identified using the genetic kinship matrix. We start by constructing a binary variable indicating whether two siblings reported different locations of birth. Assuming that the siblings grew up together, their location of birth can differ for two reasons. First, the family may have moved house between the births of their two or more children (i.e., a ‘true’ change in their birth location). Second, the location of birth may have been misreported by one or more children (i.e., an ‘error’ in their birth location). For the former, we assume that a longer spacing between births linearly increases the probability of a house move; something we test empirically below. The latter can occur due to incorrect location recall by the UKB participant, or due to differential recording by interviewers (e.g., recorded at different levels of detail). We refer to such location ‘error’ as measurement error.

We examine the relationship between differences in siblings’ reported birth location and the age gap between the siblings. This allows us to derive the probabilities of house moves as well as misreporting. We explore heterogeneity in these probabilities across a wide range of factors, including birth cohorts, district types, district population density, UKB assessment centre locations (at which the birth location was recorded), siblings’ gender, districts’ socio-economic composition, and siblings’ polygenic index (PGI) for education.

It is well-known that non-systematic measurement error in explanatory variables causes attenuation bias in the estimated coefficients in a linear regression. Measurement error in individuals’ recorded birth locations will therefore lead to attenuation bias when using the birth location or external data merged based on the location as explanatory variables. We explore the extent of this attenuation bias using Monte Carlo simulations for measures of disease exposure, demographic variables, and simulated spatial data with varying levels of spatial correlation and time variation.

## Results

### Differences in siblings’ birth location

We start by graphically presenting the unadjusted relationship between the discordance of siblings’ birth locations and their age gap in Figure 1. Panels (a)–(i) show the relationship for different levels of birth location accuracy. Panels (a)–(c) plot the share of sibling pairs reporting different parishes, districts, or counties of birth, respectively. This shows that 28–30% of twins (i.e., siblings with an age gap of 0 years) report coordinates in different parishes and districts, with 8% reporting birth coordinates that are located in different counties. Furthermore, the graphs show a clear increase in this discordance as the age gap between siblings increases. This is expected, since an increase in birth spacing also increases the likelihood of a house move between the two births.

**Figure 1:**
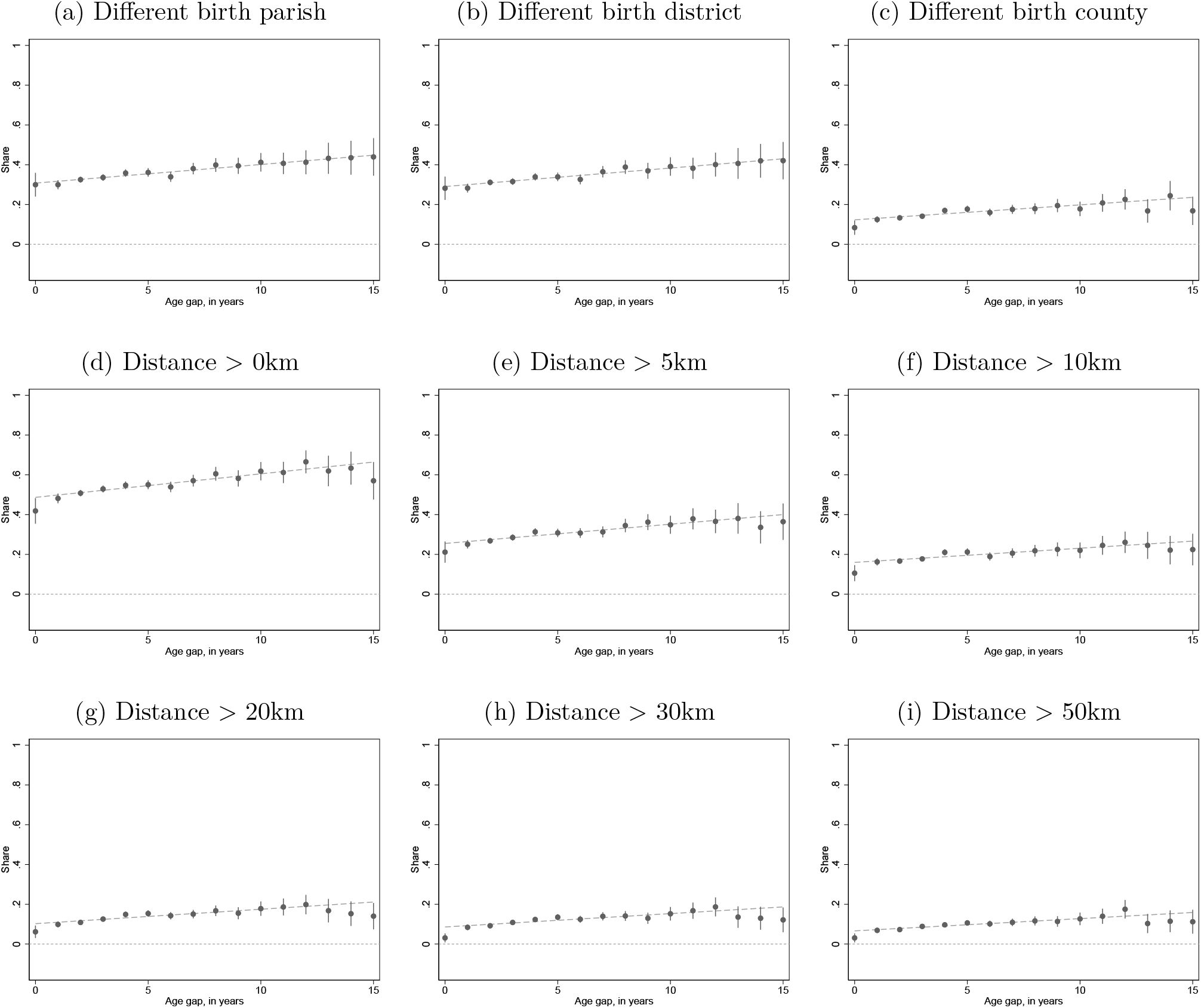
Differences in siblings’ birth location and their age gap

To explore what may be driving the relatively large proportion reporting a different geographic area of birth for those born within a small age gap, we examine the shares of siblings reporting birth locations more than 0, 5, 10, 20, 30 and 50 kilometres apart in panels (d)–(i). This shows a similar positive relationship between the age gap and the probability of siblings reporting different coordinates. In fact, we show below that the slope coefficient in a linear regression is very similar across specifications. Furthermore, we show that the discordance between siblings is mainly driven by relatively small differences in eastings and northings. Indeed, 42% of twins report birth location coordinates that differ (i.e. are more than zero kilometres apart), but this reduces to 21% when we define discordance as those who report locations at least five kilometres apart, 11% at ten kilometres and 6% at twenty kilometres. From thirty kilometres onwards, the discordance share among twins is fairly stable at 3%.

We next quantify the relationship between siblings’ age gap and the discordance of their reported birth location further using a linear regression. In addition to the estimated discordance for twins (i.e., those with an age gap of zero; the constant), the top panel of Table 1 shows that each additional year between the birth of two siblings increases the probability of reporting different coordinates by approximately 1 percentage point. While this estimate is very similar across the different specifications, it does decrease with distance, indicating that the probability of a “long-distance move” is lower than the probability of any move.

**Table 1:** Differences in siblings’ birth location and their age gap.

|  | Different birth location: |  |  |  |  |  |  |  |
| --- | --- | --- | --- | --- | --- | --- | --- | --- |
|  | (1)<br>Parish | (2)<br>District | (3)<br>County | (4)<br>d>0km | (5)<br>d>5km | (6)<br>d>10km | (7)<br>d>30km | (8)<br>d>50km |
| Age gap (years) | 0.009***<br>(0.001) | 0.009***<br>(0.001) | 0.008***<br>(0.001) | 0.012***<br>(0.001) | 0.010***<br>(0.001) | 0.007***<br>(0.001) | 0.007***<br>(0.001) | 0.006***<br>(0.001) |
| Constant | 0.308***<br>(0.006) | 0.290***<br>(0.006) | 0.123***<br>(0.005) | 0.487***<br>(0.006) | 0.255***<br>(0.006) | 0.160***<br>(0.005) | 0.086***<br>(0.004) | 0.066***<br>(0.004) |
| **Derived probabilities:** |  |  |  |  |  |  |  |  |
| $\hat{q}$ (move probability) | 0.009***<br>(0.001) | 0.009***<br>(0.001) | 0.008***<br>(0.001) | 0.012***<br>(0.001) | 0.010***<br>(0.001) | 0.007***<br>(0.001) | 0.007***<br>(0.001) | 0.006***<br>(0.001) |
| $\hat{p}$ (error probability) | 0.168***<br>(0.004) | 0.158***<br>(0.004) | 0.063***<br>(0.002) | 0.284***<br>(0.004) | 0.137***<br>(0.003) | 0.083***<br>(0.003) | 0.044***<br>(0.002) | 0.034***<br>(0.002) |
| N (sibling pairs) | 18,482 | 18,482 | 18,482 | 18,482 | 18,482 | 18,482 | 18,482 | 18,482 |

Note: The sample is restricted to the first sibling pair observed in each family. Heteroskedasticity robust standard errors are shown in parentheses. Standard errors for $\hat{p}$ were computed using the delta method. Significance levels are indicated as follows: \* $p<0.1$ , \*\* $p<0.05$ , \*\*\* $p<0.01$

Finally, we investigate potential non-linearities in the relationship between the discordance of birth location and siblings’ age gap. Appendix Table B.1 shows that adding a quadratic term only marginally changes the estimated probabilities, suggesting that the linear specification in Table 1 is appropriate.

### Derived probabilities of household moves and measurement error

The regression estimates in the top panel of Table 1 can be used to derive the estimated annual probabilities of a household move 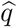 (equivalent to the slope coefficient on the sibling age gap in the top panel), as well as the probability of measurement error in the reporting of an individual’s birth locations 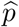 (derived using Equation 4). These derived probabilities are shown in the bottom panel of Table 1.

The probability of a household move to different geographical coordinates (at a 1 kilometre resolution; column 4) during the childhood of UKB participants is estimated to be 1.2% annually. The annual probability of moving to a different parish or district is estimated at 0.9%. As can be expected, the probability of long-distance moves is lower with an annual probability of 0.8% for moves to a different county and 0.6% for a move of more than 50 kilometres.

The probability of an error in participants’ birth coordinates (at a 1 kilometre resolution; column 4) is estimated to be 28.4%. However, a large share of these errors is due to small differences in the birth location - with the estimated error probability reducing to 13.7% and 8.3% for differences of more than 5 and 10 kilometres respectively. Similarly, the estimated probability of an error in a participant’s birth parish and district are 16.8% and 15.8% respectively. Errors with a large difference in birth location are relatively rare, with an estimated error probability of 6.3% for participants’ county of birth, and 3.4% for birth location differences of more than 50 kilometres.

In Appendix C we explore heterogeneities in the annual probability of a household move and the probability of measurement error along several dimensions. We find that later (i.e. younger) cohorts and those living in rural, less densely populated areas exhibit more measurement error. Our results furthermore show that mobility is higher among families initially living in rural and highly educated areas. Finally, we observe substantial differences in measurement error across the assessment centre locations at which participants completed their initial interview.

### Attenuation bias

Measurement error in the birth location may lead to attenuation bias when investigating the impact of birth-location-based variation in exposures. The bias is increasing in the variance of the measurement error, which is driven by (i) the probability of an error in the birth location, (ii) by the difference in the early life environment between the true and the reported place of birth, and (iii) by the share of (non-spatial) variation over time exhibited by the environmental measure. Environmental measures at a more granular, less aggregated level (e.g. at the precise coordinate level) are subject to a higher probability of error in the birth location, which will increase the measurement error and thus the attenuation bias. High levels of spatial correlation in the early life environment reduce the consequences of errors in individuals’ birth location reports, and with that the resulting attenuation bias. In contrast, low levels of spatial correlation lead to increased attenuation bias, since small errors in birth locations can have large consequences for the measurement error in the exposure variable. Variation in the early life environment over time is not affected by the measurement error of birth locations, thus the larger the share of variation in the exposure that is driven by variation over time (as opposed to spatial variation), the smaller the susceptibility to the attenuation bias.

Sibling fixed effects models may be particularly vulnerable to the attenuation bias as they rely on differences in exposure between siblings. Such differences in exposure may be due to siblings’ differences in their location or date of birth. In cases where the exposure mainly captures spatial (rather than temporal) variation, sibling differences in the exposure will be largely driven by measurement error. Based on our estimates in Table 1 and the average age gap between siblings of 4.5 years, we conclude that approximately 80-85% of birth location differences are driven by misreporting and only 15-20% are due to house moves. In other words, variation due to measurement error will dominate true variation in the measured exposure when time variation in the exposure is low. With higher levels of time variation, the share of true variation in the measure of exposure will increase and the attenuation bias will reduce.

We quantify the size of the attenuation bias using Monte Carlo simulations for a variety of exposures with varying levels of spatial and temporal variation, and for ordinary least squares (OLS) as well as sibling fixed effect specifications. Figure 2 shows the size of the attenuation bias for simulated spatial data at the district-of-birth level (we repeat this exercise at coordinate and parish levels of geographical aggregation in Appendix D). Panel a) gives the percentage attenuation bias of the slope coefficient in a bivariate linear regression. For exposure variables without spatial correlation and time variation, the bias is predicted to be approximately 16.5%. As the spatial correlation increases, the consequences of measurement error in the birth location for the exposure variable are reduced and the attenuation bias shrinks. As the share of time variation increases, a smaller share of the overall variance in the exposure is subject to measurement error and the attenuation bias is reduced. Exposures with 100% of variance due to time variation are no longer subject to any bias, as spatial measurement errors no longer have any consequences on the exposure. The maximum bias for exposures at the birth-coordinate (Figure D.1) and parish level (Figure 2) is 28% and 17% respectively, and again the bias decreases as spatial correlation and time variation of the exposure increase.

**Figure 2:**
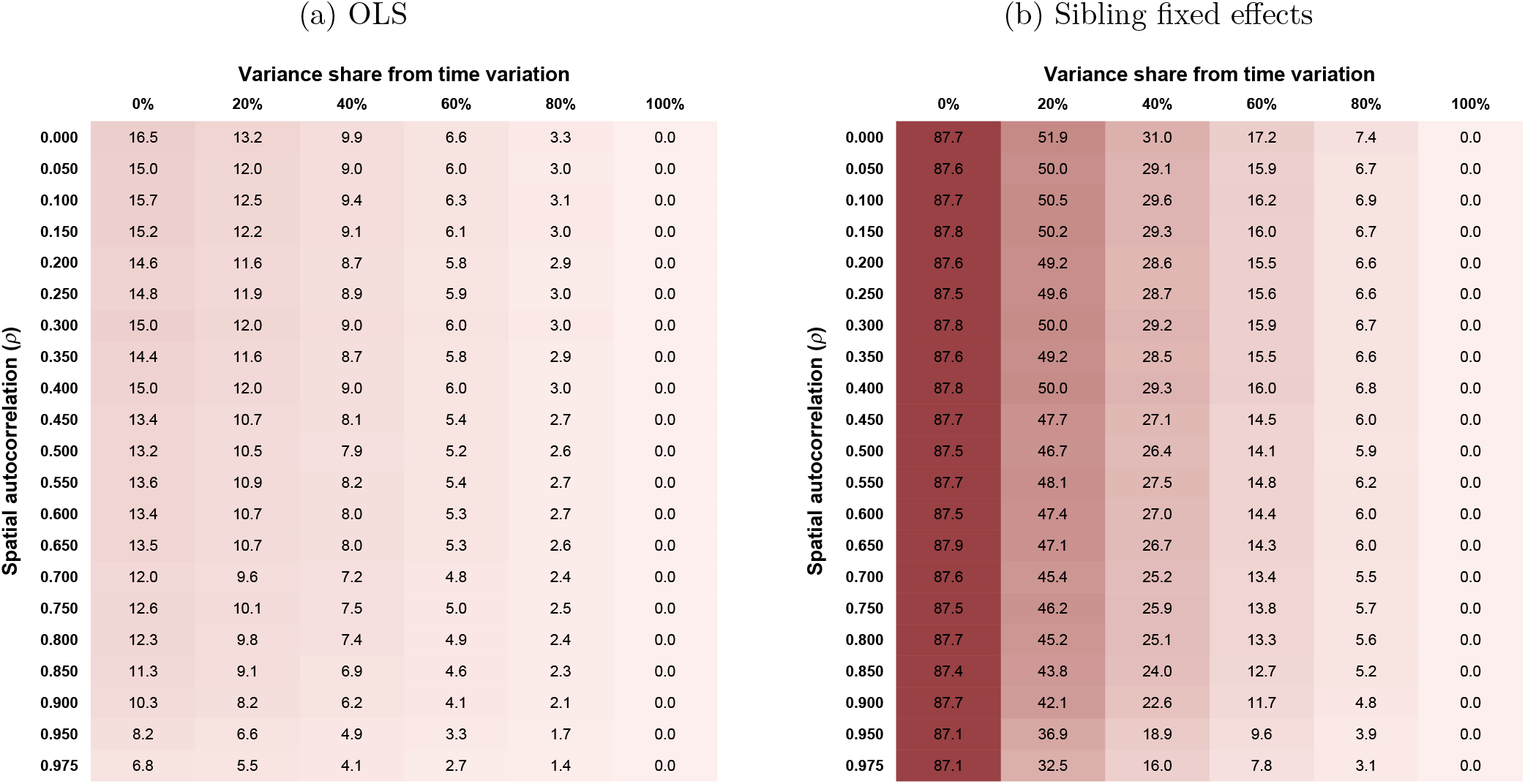
Attenuation bias for district-level data with different levels of spatial correlation and time variation Note: The attenuation bias values shown are the mean bias (in %) from simulations of OLS and sibling fixed effects estimations with 1000 repetitions. For each level of spatial autocorrelation (*ρ*), 10 district-level variables were simulated and merged to the sibling sample. The district-level spatial variables were combined with normally distributed year-month of birth fixed effects to simulate time-varying spatial exposures. The columns of the tables correspond to different ratios of spatial to temporal variation when simulating the exposure variable - as indicated by the share of the exposure variance due to time variation. Each simulated variable was then used in 100 simulations of the attenuation bias based on an error probability for the district of birth *p* = 0.158 and a move probability *q* = 0.009.

In Figure 3 we present the simulated attenuation bias for examples of previously-studied or otherwise-relevant district-level exposure variables with different levels of spatial and temporal variation which we obtain from alternative data sources (see note to figure). The predicted bias of OLS estimates ranges from 5.6% for the exposure to the infant mortality rate during the first year of life to 23% for the time-invariant share of social class III from the 1951 Census. Note, that the bias of OLS estimates for some of these examples exceeds the bias for simulated district-level exposures in Figure 2. While the simulated exposures assume identical distributions of exposures for sibling pairs reporting the same birth location and sibling pairs who do not, this does not hold for all examples and can lead to a larger attenuation bias if the variance of an exposure is larger among sibling pairs reporting different birth locations.

**Figure 3:**
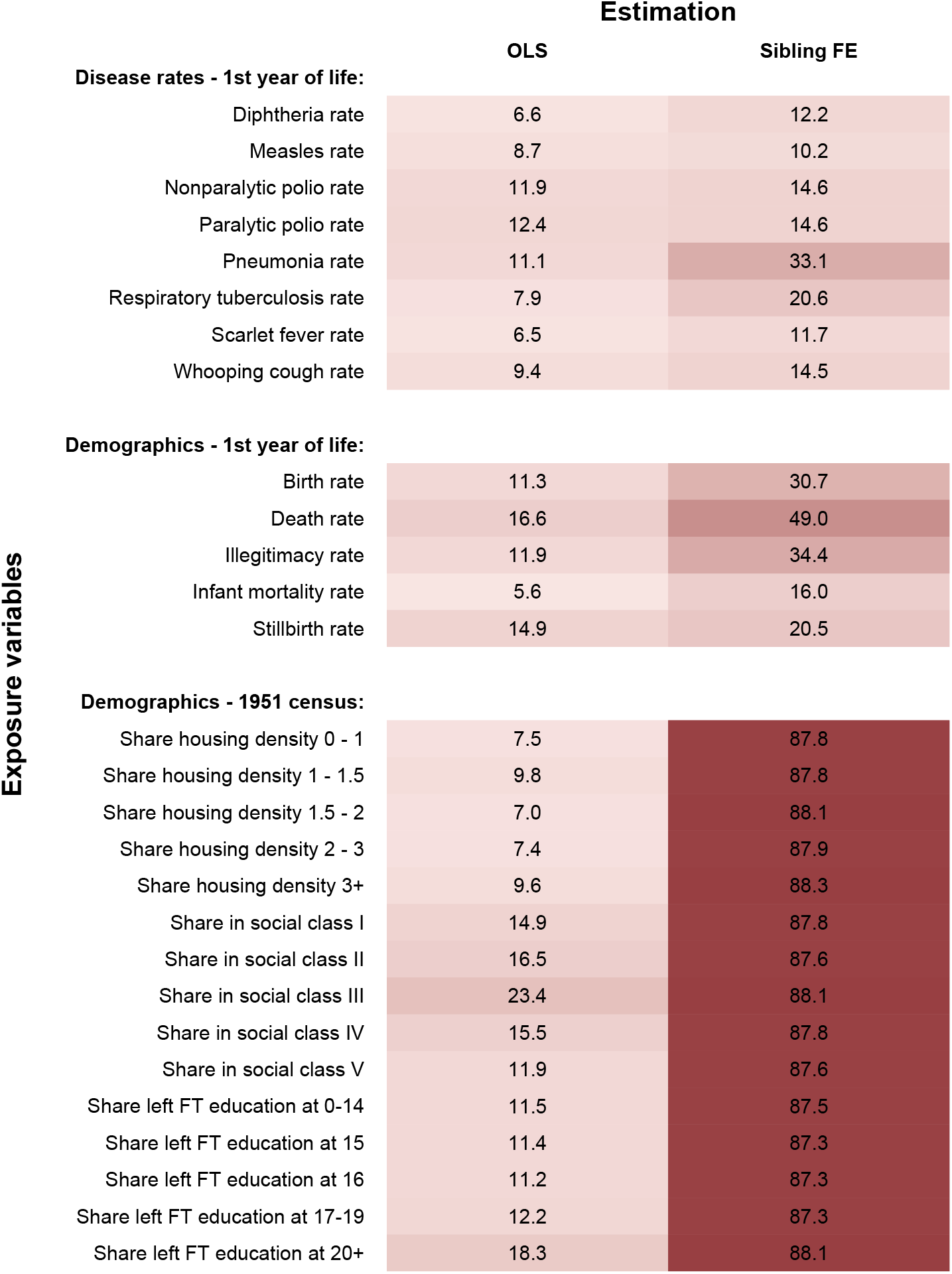
Attenuation bias for different district-level exposure variables Note: The attenuation bias values shown are the mean bias (in %) from simulations of OLS and sibling fixed effects estimations with 1000 repetitions based on an error probability for the district of birth *p* = 0.158 and a move probability *q* = 0.009. Disease data are from the Registrar General’s Weekly Reports ^22^,23, demographic data are from the Registrar General’s Statistical Review of England and Wales ^24^ and the 1951 census ^25^,26.

Panel b) of Figure 2 reports the attenuation bias in sibling fixed effect models for simulated district-level exposures. We find a very large attenuation bias of 87-88% when the exposure variable does not vary over time. For data at the birth-coordinate (Figure D.1) and parish level (Figure D.2) the bias is up to 90% and 88% respectively. Even an increase in spatial correlation does not reduce this bias substantially in the absence of any time variation, since the true variation in the exposure from household moves and the false variation from measurement error decrease at the same rate. For exposures that do vary over time and space, an increase in spatial correlation does reduce the bias. Furthermore, increases in the temporal variation reduce the bias substantially, shrinking it to zero for exposures driven solely by time variation. For the example district-level exposure variables (Figure 3), the attenuation bias in a sibling fixed effects model is substantially larger than in bivariate linear regressions. The predicted bias ranges from 10% for exposure to highly time-varying measles rates to 87-88% for the census-based demographic measures as they are (by construction) time-invariant. These findings are in line with the measurement error literature^27,28^ which shows that fixed effects estimations may aggravate the attenuation bias, especially in cases where the signal is highly correlated over time (i.e. little time variation in the true exposure) but the error is not (i.e. no or little correlation between siblings’ birth location errors).

### Consequences of discordance in siblings’ birth location for the spatial correlation of genetic principal components

Principal components of genotype data are spatially correlated within the United Kingdom^4,7^. Measurement error in birth location data and household mobility is therefore expected to affect the strength of these spatial correlations. We explore this in the following. Figure E.1 confirms the strong levels of spatial correlation for the first five principal components in the UKB (based on principal component analysis for a homogeneous population of white-British UKB respondents; Moran’s *I >* 0.8 at the district level). We find much less spatial correlation for the sixth principal component and therefore do not focus our discussion on the latter.

The vertical axis of Figure 4 shows the correlation of each genetic principal component between individuals in the sibling sample and the mean among individuals in the non-sibling sample who reported the same birth location. We estimate these correlations separately for siblings with different levels of discordance in their reported birth location, measured along the horizontal axis. Correlations between siblings without any discordance in birth location and others reporting the same birth location are above 0.35 for the first five principal components. As the distance between siblings’ reported birth location increases in Figure 4, the correlation with others reporting the same birth location reduces. Indeed, when comparing siblings who reported being born more than 200 kilometres apart (3.6% of sibling pairs) to those without any discordance the correlation coefficients decrease by more than 40% for all five spatially structured principal components. These results illustrate the impact of household mobility and measurement error on the estimation of the spatial structure of genetic data.

**Figure 4:**
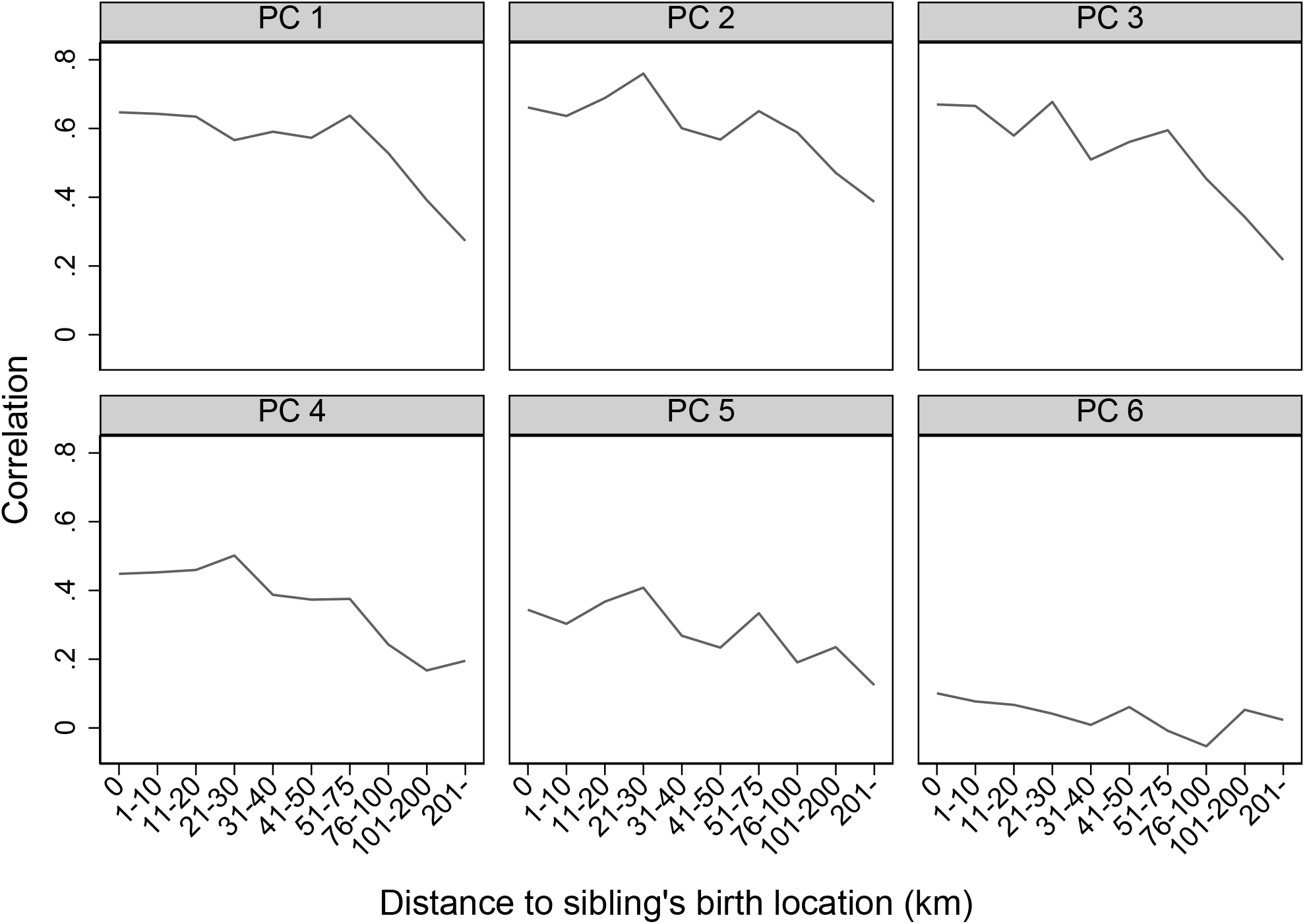
Correlation of siblings’ genetic principal components with others reporting the same birth location Note: The figures show the correlation between genetic principal components of individuals in our sibling sample with the mean genetic principal components of non-sibling UKB participants who reported the same birth coordinates. Correlations are shown separately for the first 6 genetic principal components (PC1 -PC6) and for distances between the reported birth locations of the individuals and their siblings. Principal components are based on principal component analysis conducted on unrelated white-british individuals in the UKB. SNPs were filtered based on minor allele frequency > 0.01 and clumped for linkage disequilibrium based on minor allele frequency (*R*^2^ *>* 0.1). Long-range linkage disequilibrium regions were removed.

## Discussion

Many data surveys ask participants to retrospectively record their residential location of birth. This information has been used in many research papers across a wide range of empirical applications. Despite their frequent use, there is little information on how accurate these data are, especially since they often rely on individuals’ accurate 30+ year recall. To address this, we explore the accuracy of retrospectively recorded birth location data by studying the sample of full siblings in the UK Biobank. Assuming that siblings grew up together, our analysis allows us to decompose the discordance in birth location into two components and quantify the importance of both: household moves and measurement error. Our estimates suggest that household mobility during the early childhood period was low, with an estimated average annual move probability of 1.2%. Although our results show substantial measurement error in participants’ birth location, in the majority of cases, participants report nearby locations, meaning that most errors are of a short distance.

The consequences of this measurement error therefore depend on what the birth location data are used for. We focus on the attenuation bias that results from the use of external data merged based on individuals’ location and date of birth as explanatory variables in a regression. We quantify the size of this bias using Monte Carlo simulations for a variety of exposure variables with varying levels of spatial correlation and time variation, including disease exposures and demographic variables.

We show that, as the share of temporal variation as well as the spatial correlation in the variable of interest increases, attenuation bias reduces. Hence, the estimated impacts of exposures that show substantial temporal variation, such as disease or infant mortality rates, are substantially less attenuated compared to the impact of exposures that are time-invariant, such as measures of socio-economic composition. We show that sibling fixed effect specifications, only exploiting variation in the outcomes and variables of interest *within* families (between siblings), are particularly vulnerable to the attenuation bias, with the bias exceeding 80% when using time-invariant exposure variables. We thus recommend caution when using the UKB birth location data, especially when estimating sibling fixed effects models where the variable of interest has limited time variation and thus a low ratio of signal to error.

For analyses which do not focus on within-sibling variation, a potential sensitivity analysis is to limit the sample to siblings which reported the same birth location. While this means omitting non-siblings as well as siblings which experienced house moves in childhood from the analysis, it allows focusing on a sub-sample with more reliable (i.e., with less error in the) birth location data. While it may be tempting to use siblings’ reported birth location as an instrument for the imperfectly measured birth location of individuals to address the measurement error and resulting attenuation bias, we do not recommend using such an Obviously Related Instrumental Variables (ORIV) approach^29^ in this case. Due to the measurement error not being classical, ORIV estimates are unlikely to overcome the bias^28^ and indeed simulations show that ORIV estimates remain biased in our setting.

A common application of sibling fixed effects models are studies of gene-environment interactions, since the random within-sibling variation in genes allows a causal interpretation^30^. For these types of analyses, we recommend using an alternative approach if the environmental measure (based on birth date and place) is exogenous both within and between siblings. In these cases, it is better to use the deviation of a sibling’s genetic measure from the mean of the sibling pair (or group) as an exogenous source of genetic variation in the sibling sample^14,21^, but use both within- and between-sibling variation in the environmental measure to avoid attenuation bias from measurement error in the birth location. Instead of using this sibling mean deviation, one can alternatively control for the imputed parental genotypes^31^.

More generally, our results highlight the impact of measurement error in birth location data on regression estimates that exploit this information. We show substantial attenuation bias that is a function of the spatial correlation and temporal variation of the variable of interest. Finally, we show that the measurement error also impacts on the spatial structure of the genetic data.

## Methods

### UK Biobank sibling sample

The UK Biobank (UKB) is a large-scale, mostly biomedical database of over 500,000 individuals living in the UK. We focus on the sample of full siblings, identified using their genetic data, which includes 41,451 UKB participants. The UKB did not explicitly sample families or households, but nevertheless does contain a substantial number of related individuals. Family relationships to other participants were not recorded in the interviews, but we can identify biological relatives based on their genetic relatedness. By using the kinship matrix provided by the UKB ^32^, we can identify related participants and derive their relationship. The kinship matrix contains relatives up to the third degree and was constructed using the KING software^33^. A kinship coefficient of approximately 0.25 (interval: 0.1770 - 0.3540) suggests that the individuals are either a parent-offspring pair or full siblings. One can then distinguish between these two types of relationships by using the identity by state (*IBS*_0_) coefficient: a coefficient above 0.0012 suggests that the pair of individuals are siblings rather than parent and offspring.

The main variables of interest in our analysis are individuals’ east and north coordinates of birth (field IDs 129 and 130), reported at a one kilometre resolution. We restrict the sample to families with at least two siblings born in England, Wales and Scotland (3,392 participants dropped) and use the birth coordinates to identify their parish, local government district (henceforth: district), and county of birth, using the regional boundaries in 1951^26,34^. Keeping the boundaries fixed over time ensures that any regional boundary changes cannot drive any differences in the (e.g.) parish or district of birth. Our sibling sample covers 2,518 parishes, 1,363 districts and 98 counties.

Additionally, we restrict the sample to the oldest two siblings observed in each family (1,095 participants dropped). Our final sample comprises 36,964 siblings from 18,482 families. Appendix Table A.1 presents some descriptive statistics on our sibling sample, showing that the age gap between siblings varies between 0 and 27 years, with a mean of 4.5 years. The data comprises 227 pairs of twins, and 57.7% of the sibling sample are female. On average, siblings report being born 22 kilometres apart, though this is highly skewed, ranging from 0–1060 km. The sibling sample is relatively similar in individual and district characteristics compared to the full UKB sample (see Appendix Table A.2), with 63–66% having an upper secondary qualification and 84–86% born in an urban or municipal district.

### Polygenic indices

We explore heterogeneity of our results with respect to the siblings’ genetic predisposition for educational attainment. For this purpose, we use the polygenic index (PGI) for education provided in the PGI repository ^35^ to split the sample into quartiles and below-/above-median sub-samples. Specifically, we use the single-trait PGI provided in the repository for what they call the first partition of the UKB which includes all siblings. This PGI is based on a discovery sample of *N* = 984, 323 which includes the other two partitions of the UKB as well as other datasets such as 23andMe, AddHealth and HRS. In line with the current genetics literature, we restrict our sample in estimations involving PGIs to those of European ancestry. Thus the sample size reduces from 18, 482 sibling pairs in our main estimations to 18, 048 sibling pairs in estimations with the PGI. In this sample, the PGI for educational attainment has an incremental *R*^2^ of 10.9%.

### Comparison of siblings’ birth locations

We start by constructing a binary variable indicating whether the siblings reported being born in a different location. The variable *y*_*s*_ equals one if either the north or east coordinates differ between the two full siblings of family *s*. We then estimate the following linear probability model:

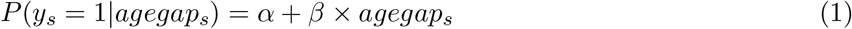

where *agegap*_*s*_ is the age gap (in years) between the two siblings in family *s*. Hence, this assumes that the discordance in reported birth location is driven by two processes: (1) ‘true’ differences due to house moves and (2) reporting/measurement error. The extent to which *agegap*_*s*_ can explain variation in *y*_*s*_ captures the former; the intercept *α* captures the latter. We explore potential non-linearities in this relationship in Appendix B. We furthermore estimate linear probability models as described by Equation 1 for a variety of alternative definitions of *y*_*s*_: different parishes of birth, different districts of birth, different counties of birth, different birth locations that are more than 5 / 10 / 20 / 30 / 50 kilometres apart.

Birth date variables in the UKB are recorded at the year-month level rather than at the daily level. This rounding introduces some classical measurement error to the age gap variable used in our estimations. The expected attenuation bias from this measurement error can be derived to be 0.01% and thus should not have any substantial impact on our estimates. Further measurement error in the reported birth date variable (and thus in the age gap between siblings) is likely to be small. Suggestive evidence of the reliability of the birth date and age gap variable is that we do not observe any sibling pairs with an age gap of 1 to 8 months, as would be expected based on the length of human gestation.

### Deriving probabilities of household moves and measurement error

We use the regression estimates of Equation 1 to derive the probability of a household move during childhood as well as the probability of measurement error in the location of birth. To derive these probabilities, we make the following assumptions:

#### Assumption 1

*If both siblings in a sibling pair report the same birth location, we assume this is the true birth location for both*.

Assumption 1 recognises that we cannot identify misreporting of birth locations if both siblings report the same *incorrect* birth locations. Ignoring these unlikely cases will downward bias the derived error probability. We furthermore cannot identify household moves in cases where the siblings incorrectly report the same birth location.

#### Assumption 2

*We assume that biological siblings grew up in the same household*.

Assumption 2 ignores adoptions shortly after birth and other events that may result in biological siblings growing up in different households. However, the frequency of such events in the UK during 1940-70 was likely low and thus any resulting upward bias in the derived error probability will be small.

#### Assumption 3

*We assume that the probability of a move between the birth of two siblings is a linear function of the age gap between them*.

We empirically test Assumption 3 in Appendix B by allowing for non-linearities in the relationship between the sibling age gap and the probability of house moves. While we find minor non-linearities, these only affect the derived error probabilities to a small degree.

#### Assumption 4

*We assume that measurement error occurs randomly with the same probability for any participant, and that these errors are independent within sibling pairs and independent of the age gap between siblings*.

Assumption 4 is unlikely to hold in reality. Indeed, we show in Appendix C that certain participant characteristics affect the probability of house moves and measurement errors. It is furthermore likely that error occurrence is positively correlated among siblings, but our data does not allow us to quantify this. Ignoring such a positive within-sibling correlation will downward bias the derived error probability. However, simulations show that a very strong sibling correlation is required to create substantial bias.

Under the above assumptions, the probability of a difference in the reported birth location between two siblings can be written as:

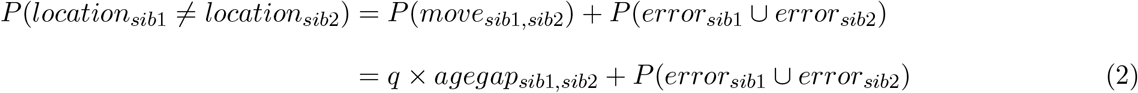

where *q* denotes the annual probability of a household move to a different location. The probability of an error in the birth location of either sibling can be written as:

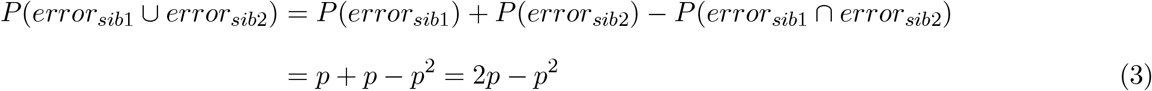

We use this to derive the probability *p* of an error in the birth location of any respondent, defined as:

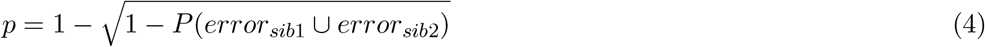

### Attenuation bias

Measurement error in the birth location data may have consequences for the use of birth-location-based explanatory variables in regression analyses. “Classical” measurement error in explanatory variables causes coefficient estimates in a linear regression to be biased towards zero^28,36^. Our application is unlikely to show classical measurement error. In fact, one would expect a negative correlation between the true explanatory variable and the measurement error. If an exposure is high (low) for the true birth location, it is more likely that the exposure in the misreported birth location is below (above) the true exposure level due to regression to the mean. Nevertheless, errors in the birth location data may lead to attenuation bias when the birth location or external location-based data are used as explanatory variables. In the following, we use the case of classical measurement error as an illustration of the attenuation bias. Consider the bivariate linear model:

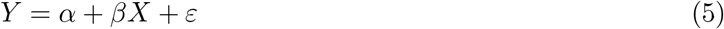

where *Y* denotes the outcome of interest, *X* is individuals’ early life environment, and *ε* is an idiosyncratic error term. Since we cannot observe *X* directly, we use a measure based on individuals’ reported date and place of birth. This measure of early life environment *X*^*\**^ is subject to measurement error *u* due to reporting / recording errors of the birth place as well as due to household moves during childhood:

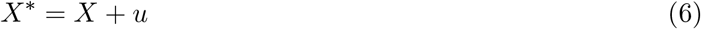

Estimating Equation 5 using the observable measure *X*^*\**^ instead of *X* leads to attenuation bias in 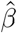 due to the measurement error:

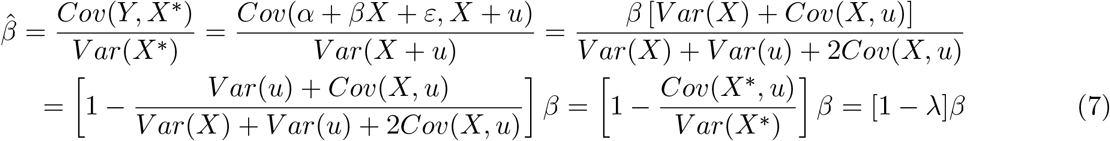

where *λ* is the magnitude of the attenuation bias. *λ* is increasing in the variance of the measurement error, which is driven by (i) the probability of an error in the birth location, (ii) by the difference in the early life environment between the true and the reported place of birth (i.e., *X* and *X*^*\**^), and (iii) by the share of (non-spatial) variation over time exhibited by the environmental measure as this temporal variation is not subject to the measurement error.

We use Monte Carlo simulations to quantify the size of the attenuation bias in ordinary least squares (OLS) and sibling fixed effect estimations for simulated data at the coordinate, parish and district level with varying levels of spatial and temporal variation, as well as for several district-level measures of disease exposure and demographics.

#### Simulation of time-varying spatially correlated data

We begin by simulating time-invariant spatially correlated data in R using the sim_sar command of the geostan package (for parish- and district-level data) and the powerWeights command of the spatialreg package (for coordinate-level data). We simulate 10 variables *S*_*a,ρ*_ for each spatial aggregation level *a* (coordinate-, parish- and district-level) and spatial autocorrelation parameter *ρ ∈* [0.00, 0.05, 0.10, …, 0.90, 0.95, 0.975].

To add time-variation to the spatial data, we draw year-month of birth fixed effects from a standard normal distribution (without any temporal autocorrelation). We simulate a year-month of birth fixed effect variable *T* for each of the simulated spatially correlated time-invariant variables *S*_*a,ρ*_. Finally, we create time-varying spatially correlated variables for different shares of variance due to temporal variation *k ∈* [0.0, 0.2, 0.4, 0.6, 0.8, 1.0] by combining the normalized time-invariant spatial variables and the normalized birth-date fixed effects as follows:

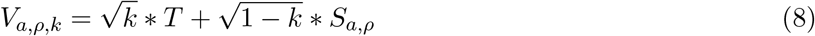

We normalize the resulting time-varying spatially correlated *V*_*a,ρ,k*_.

#### Simulation of attenuation bias

The simulated time-varying spatially correlated variables *V*_*a,ρ,k*_ as well as a variety of district-level measures of disease exposure and demographics are merged with the sibling sample used in our main estimations based on each individual’s reported birth date and birth location; the resulting variable is defined as *X*_*t,ownloc*_. Additionally we also merge these variables based on each individual’s reported birth date and their sibling’s reported birth location (*X*_*t,sibloc*_), as well as the geographic midpoint between the two birth locations reported by the two siblings (*X*_*t,midloc*_). If the geographic midpoint between the two birth locations is not on land, the closest UK land location to the midpoint is used.

For each sibling pair with different birth locations (defined at the coordinate, parish or district level) we compute the predicted probability of an error in the reported birth location conditional on different locations being reported:

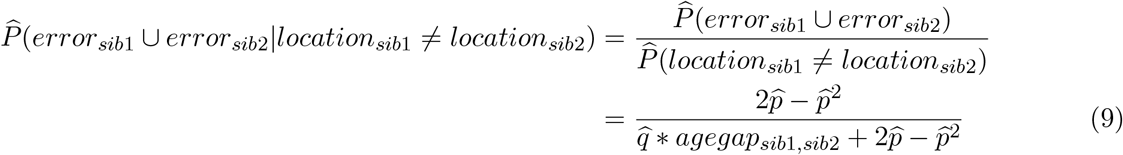

based on the estimates 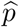 and 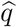 from the main analysis (Table 1) and the siblings’ age gap.

For each variable we then simulate the attenuation bias using 1000 repetitions. For the simulated time-varying spatially correlated variables *V*_*a,ρ,k*_, we run 100 repetitions for each variable. With 10 variables for each level of aggregation *a*, autocorrelation *ρ* and temporal variance share *k*, this corresponds to 1000 simulations. Each repetition of the Monte Carlo simulation proceeds as follows: For sibling pairs with different reported birth coordinates / districts / parishes, we draw a Bernoulli random variable *E*_*s*_ indicating whether an error occurred (*E*_*s*_ = 1) using the error probability 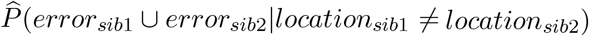 computed above. For sibling pairs with the same reported birth coordinates / district / parish, we set *E*_*s*_ = 0. A second Bernoulli variable *B*_*s*_ is then drawn for those sibling pairs with *E*_*s*_ = 1 to indicate whether both sibling birth locations are subject to an error, based on the conditional probability 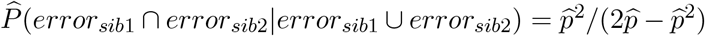. If only one of the birth locations is incorrect (*E*_*s*_ = 1, *B*_*s*_ = 0), then one of the two siblings is chosen at random for the error.

The “true” exposure in our simulations is defined as follows:

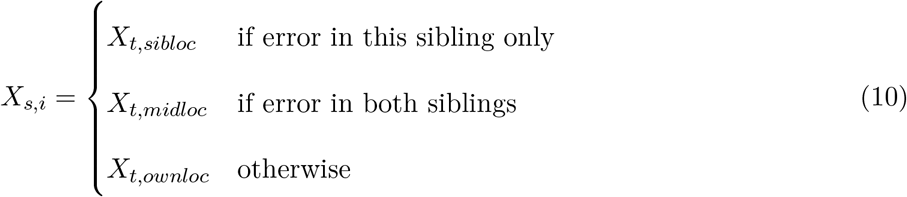

Irrespective of any errors to the reported birth location, the exposure *X*_*s,i*_ is always defined based on the individual’s reported date of birth *t*. If only one sibling in a sibling pair is simulated to have an incorrect birth location, their sibling’s reported birth location *sibloc* is used to compute the exposure. If both siblings in a sibling pair are simulated to have an incorrect birth location, we use the mid-point *midloc* between their reported birth locations. In the absence of any information on the true birth location in these cases, the midpoint between the two reported locations sets a lower bound on the average geographic distance between the true and the observed birth locations. For all individuals who are not simulated to have an incorrect birth location, we use their own reported birth location *ownloc*.

The observed exposure in our simulations is defined as

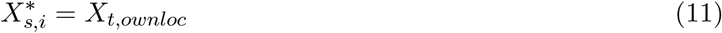

for all individuals and thus subject to measurement error due to misreported birth locations.

We simulate an outcome *Y*_*s,i*_ which is affected by the “true” exposure as follows:

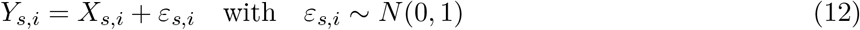

We then calculate the OLS attenuation bias by comparing the coefficients estimated for the following two OLS equations:

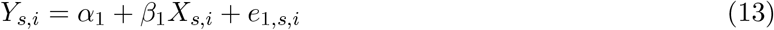

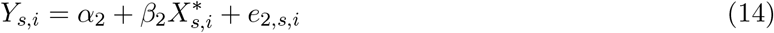

The difference between 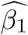 in the estimations using the “true” exposure and 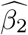 in the estimations using the observed exposure is the OLS attenuation bias. Similarly, we calculate the attenuation bias in sibling fixed effects estimations by comparing the coefficient estimates 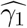 and 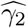 for the following two equations:

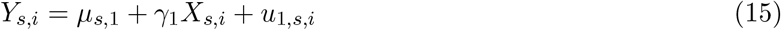

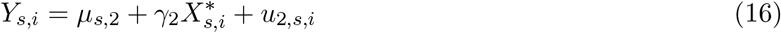

### Principal component analysis

We examine the impact of measurement error in the reported birth locations and of household mobility on the strength of spatial correlations in genetic principal components. For these analyses we conduct a principal component analyses using the big_randomSVD command of the bigstatsr package. We restrict the sample to unrelated white-British individuals in the UKB and remove genetic outliers ^37^, resulting in a sample size of *N* = 276, 279. SNPs are filtered based on minor allele frequency > 0.01 and clumped for linkage disequilibrium based on minor allele frequency (*R*^2^ *>* 0.1). Long-range linkage disequilibrium regions are removed. This results in a set of 108, 251 SNPs used in the principal component analysis. We predict the resulting principal component vectors in the estimation sample of unrelated white-British individuals described above, as well as in the sibling sample (*N* = 33, 721 after removing any genetic outliers).

## Data availability

UK Biobank data are available following an application procedure described at https://www.ukbiobank.ac.uk/enable-your-research.

## Acknowledgements

We gratefully acknowledge funding of this project from the European Research Council (ERC) under the European Union’s Horizon 2020 research and innovation programme, grant agreement no. 851725. We thank the GEIGHEI project team and participants at the European Social Science Genomics Network Conference for their helpful comments.

## Appendix For Online Publication

## Appendix A Descriptive statistics

**Table A.1:**
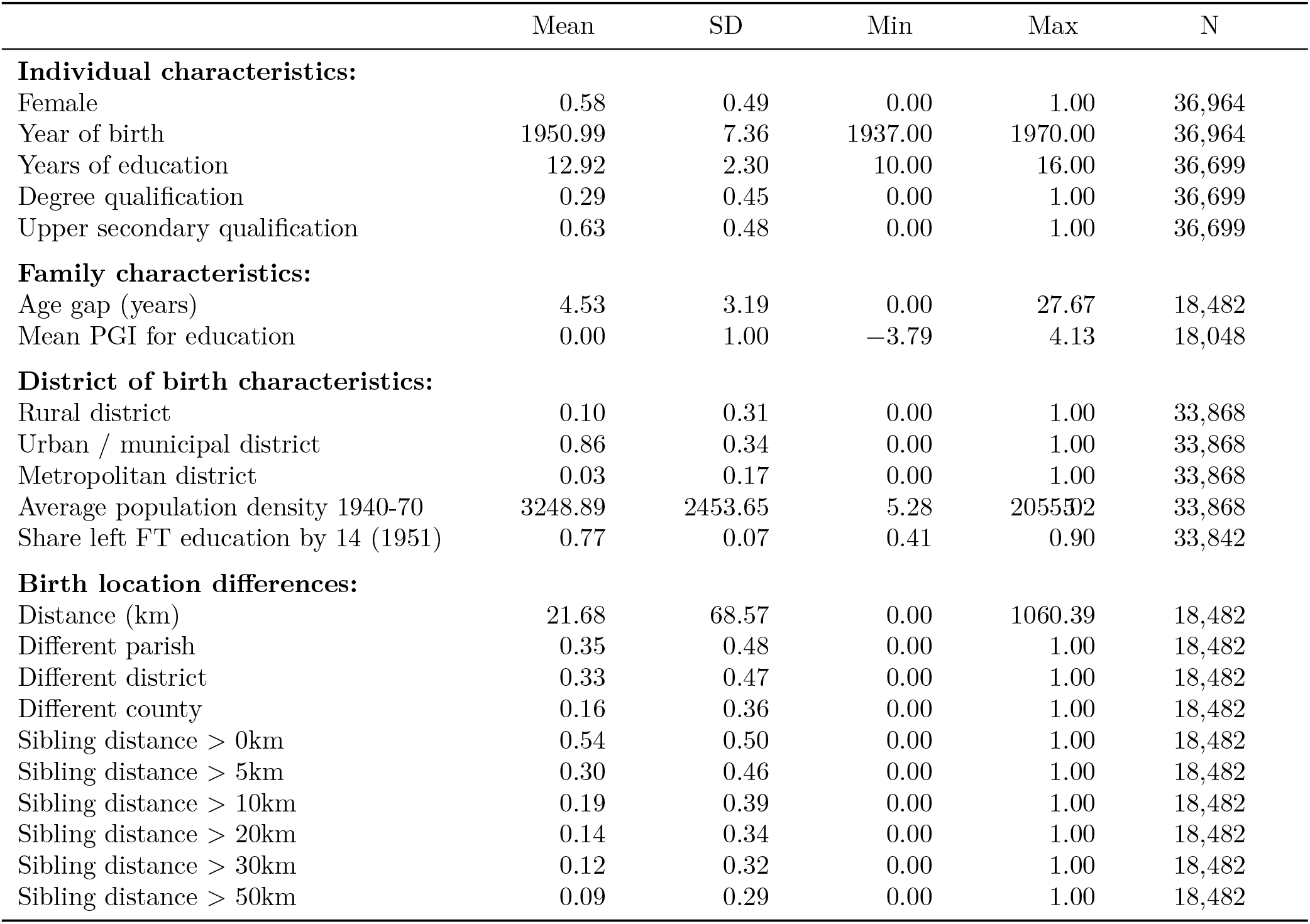
Descriptive statistics.

**Table A.2:** Comparison of sibling sample and full UK Biobank sample.

|  | Sibling sample |  |  | Full UKB sample |  |  |
| --- | --- | --- | --- | --- | --- | --- |
|  | Mean | SD | N | Mean | SD | N |
| **Individual characteristics:** |  |  |  |  |  |  |
| Female | 0.58 | 0.49 | 36,964 | 0.54 | 0.50 | 502,412 |
| Year of birth | 1950.99 | 7.36 | 36,964 | 1951.54 | 8.12 | 502,412 |
| Years of education | 12.92 | 2.30 | 36,699 | 13.11 | 2.33 | 493,364 |
| Degree qualification | 0.29 | 0.45 | 36,699 | 0.33 | 0.47 | 493,364 |
| Upper secondary qualification | 0.63 | 0.48 | 36,699 | 0.66 | 0.47 | 493,364 |
| **District of birth characteristics:** |  |  |  |  |  |  |
| Rural district | 0.10 | 0.31 | 33,868 | 0.11 | 0.32 | 405,144 |
| Urban / municipal district | 0.86 | 0.34 | 33,868 | 0.84 | 0.37 | 405,144 |
| Metropolitan district | 0.03 | 0.17 | 33,868 | 0.05 | 0.22 | 405,144 |
| Average population density 1940-70 | 3248.89 | 2453.65 | 33,868 | 3220.22 | 2752.28 | 405,144 |
| Share left FT education by 14 (1951) | 0.77 | 0.07 | 33,842 | 0.76 | 0.08 | 404,746 |

## Appendix B Sensitivity analysis

**Table B.1:** Differences in siblings’ birth location and their age gap -nonlinear specification.

|  | Different birth location: |  |  |  |  |  |  |  |
| --- | --- | --- | --- | --- | --- | --- | --- | --- |
|  | (1)<br>Parish | (2)<br>District | (3)<br>County | (4)<br>d>0km | (5)<br>d>5km | (6)<br>d>10km | (7)<br>d>30km | (8)<br>d>50km |
| Age gap (years) | 0.0152***<br>(0.003) | 0.0140***<br>(0.003) | 0.0119***<br>(0.003) | 0.0193***<br>(0.003) | 0.0181***<br>(0.003) | 0.0133***<br>(0.003) | 0.0138***<br>(0.002) | 0.0102***<br>(0.002) |
| Age gap squared | -00004*<br>(0.000) | -00003<br>(0.000) | -00003*<br>(0.000) | -00005**<br>(0.000) | -00006***<br>(0.000) | -00004**<br>(0.000) | -00005***<br>(0.000) | -00003*<br>(0.000) |
| Constant | 0.2942***<br>(0.009) | 0.2794***<br>(0.009) | 0.1128***<br>(0.007) | 0.4696***<br>(0.010) | 0.2355***<br>(0.009) | 0.1454***<br>(0.008) | 0.0694***<br>(0.006) | 0.0568***<br>(0.006) |
| **Derived probabilities:** |  |  |  |  |  |  |  |  |
| $\hat{q}(\text{agegap} = 1)$ | 0.0147***<br>(0.003) | 0.0137***<br>(0.003) | 0.0116***<br>(0.002) | 0.0188***<br>(0.003) | 0.0175***<br>(0.003) | 0.0129***<br>(0.003) | 0.0133***<br>(0.002) | 0.0099***<br>(0.002) |
| $\hat{q}(\text{agegap} = 3)$ | 0.0139***<br>(0.003) | 0.0130***<br>(0.003) | 0.0110***<br>(0.002) | 0.0177***<br>(0.003) | 0.0163***<br>(0.003) | 0.0120***<br>(0.002) | 0.0123***<br>(0.002) | 0.0093***<br>(0.002) |
| $\hat{q}(\text{agegap} = 6)$ | 0.0126***<br>(0.002) | 0.0120***<br>(0.002) | 0.0100***<br>(0.002) | 0.0161***<br>(0.002) | 0.0145***<br>(0.002) | 0.0106***<br>(0.002) | 0.0107***<br>(0.001) | 0.0085***<br>(0.001) |
| $\hat{q}(\text{agegap} = 9)$ | 0.0114***<br>(0.002) | 0.0110***<br>(0.002) | 0.0091***<br>(0.001) | 0.0144***<br>(0.002) | 0.0126***<br>(0.001) | 0.0093***<br>(0.001) | 0.0092***<br>(0.001) | 0.0076***<br>(0.001) |
| $\hat{p}$ | 0.1599***<br>(0.005) | 0.1511***<br>(0.005) | 0.0581***<br>(0.004) | 0.2717***<br>(0.007) | 0.1256***<br>(0.005) | 0.0756***<br>(0.004) | 0.0353***<br>(0.003) | 0.0288***<br>(0.003) |
| N (sibling pairs) | 18,482 | 18,482 | 18,482 | 18,482 | 18,482 | 18,482 | 18,482 | 18,482 |

Note: $\hat{q}$ is the estimated increase in the probability of at least one house move when increasing the age gap between siblings by 1 additional year, it is given above for different age gap levels. $\hat{p}$ is the estimated probability of an error in the recorded birth location of a UKB participant. The sample is restricted to the first sibling pair observed in each family. Heteroskedasticity robust standard errors are shown in parentheses. Standard errors for $\hat{q}$ and $\hat{p}$ were computed using the delta method. Significance levels are indicated as follows: \* p<0.1, \*\* p<0.05, \*\*\* p<0.01

## Appendix C Heterogeneity analysis

In this Appendix we explore heterogeneities in the annual probability of a household move and the probability of measurement error along several dimensions. Table C.1 shows limited birth cohort differences in the probability of household moves to a different district, but a larger probability of measurement error among later (i.e. younger) cohorts. The gender composition of the sibling pairs is, as expected, not found to matter for the probabilities of household moves and measurement error (Table C.2).

Table C.3 shows a substantially higher probability of moving out of rural districts at 2.1% annually, compared to 0.8% and 0.9% for urban/municipal and metropolitan districts respectively. A cross-tabulation of district types reported by the 1st (i.e. older) and 2nd (i.e. younger) sibling in Table C.4 suggests that a large share of these moves originating in rural districts was to urban / municipal districts. The probability of measurement error also differs substantially by district type, with Table C.3 showing this is 13.8% for sibling pairs with the older sibling born in urban / municipal districts, 28.2% for rural districts and 39.4% for metropolitan districts.

Table C.5 provides additional evidence based on an alternative approximation of urbanicity, comparing population density quartiles of the older sibling’s district of birth. Again, we find a higher probability of moving out of more rural areas, with an annual probability of 1.4% for the lowest population density quartile compared to 0.6% for the highest density quartile. The probability of measurement error is also found to be largest in less densely populated areas.

In Table C.6 we examine heterogeneity between areas with high and low levels of educational attainment, according to data from the 1951 census^25,26^. Mobility is higher among families living in more highly educated areas (i.e., districts with a low share of adults who left education by age 14), with an annual move probability of 1.4% in districts with the highest levels of education compared to 0.7% in districts with the lowest education levels. For the probability of measurement error, the picture is less clear but suggestive of higher error probabilities in more highly educated areas.

In Table C.7, we use the average polygenic index (PGI) for education of the two siblings to proxy for the parental PGI and examine whether mobility and misreporting are heterogeneous between families with higher and lower polygenic indices for education. The PGI is the best linear genetic predictor of education, but does not necessarily capture biological or non-modifiable effects. There is no systematic heterogeneity in the move probability, but some suggestion – perhaps unexpectedly – that error probabilities are slightly higher among families with a higher genetic propensity for education.

Previous research shows selective migration based on one’s PGI^4^. More specifically, they show that individuals who leave UK coal mining areas carry more education-increasing alleles compared to individuals elsewhere in the UK, highlighting genetic correlates of social stratification. We explore this in an alternative way, examining whether a family’s genetic propensity for education differentially affects mobility in areas with low and high levels of education. Table C.8 indeed shows a slightly higher probability of moving for those with above-median PGIs, but in contrast to previous research, we find larger move probabilities in higher educated areas. This contrast in findings may be explained by the fact that we are examining mobility in the 1940-1970s among families with young children, while previous findings^4^ likely capture mobility at a later time (e.g. following the closure of UK coal mines in the 1980s) during early adulthood of the UKB participants.

In addition to characteristics of the UKB participants and their area of birth, measurement error may also be related to the UKB assessment centre at which they were first interviewed. The birth location data is based on an interaction between the participant and an interviewer, who identified the place of birth from a long list of place names in the computer system based on the information given by the participant. Thus the accuracy of the recorded birth location may be subject to interviewer effects. In the absence of identifiers for the interviewer, we explore the importance of interviewer effects by studying heterogeneity in the error probability across the 22 assessment centres of the UKB. Panel (a) of Figure C.1 shows substantial heterogeneity in the discordance share for siblings’ district of birth: only 26% of participants interviewed in Edinburgh are recorded to have a different district of birth to their sibling, whilst this is 55% of those interviewed in Barts. Naturally, these differences translate to substantial heterogeneity in the estimated probability of measurement error shown in panel (b), from 11% in Edinburgh to 30% in Barts. These results suggest substantial interviewer effects in the accuracy of the birth location data.

**Table C.1:**
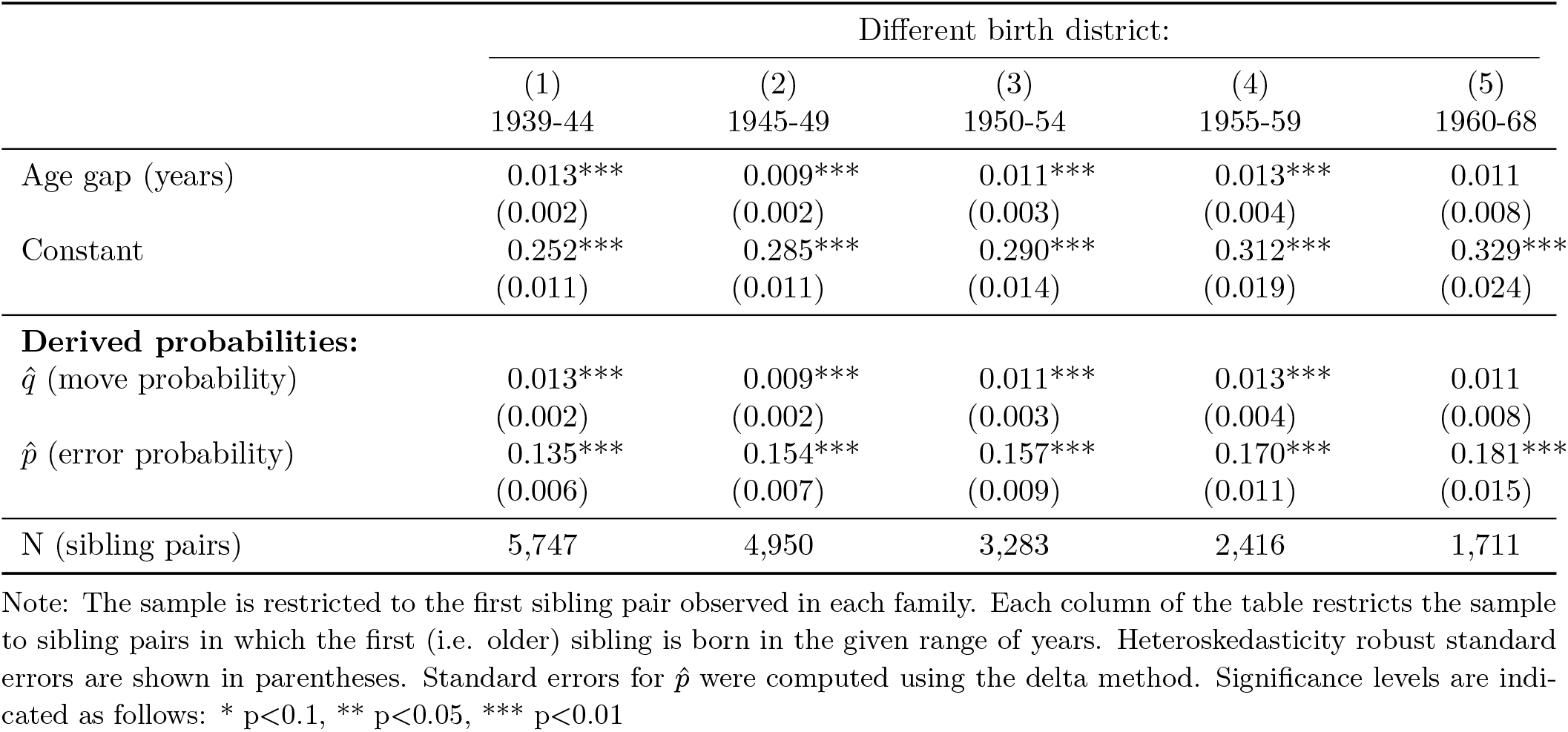
Differences in siblings’ birth district and their age gap - across different birth cohorts.

**Table C.2:** Differences in siblings’ birth district and their age gap - across sibling gender compositions.

|  | Gender composition of sibling pair: |  |  |
| --- | --- | --- | --- |
|  | (1)<br>F/F | (2)<br>F/M | (3)<br>M/M |
| Age gap (years) | 0.009***<br>(0.002) | 0.010***<br>(0.002) | 0.009***<br>(0.002) |
| Constant | 0.282***<br>(0.010) | 0.297***<br>(0.009) | 0.290***<br>(0.014) |
| **Derived probabilities:** |  |  |  |
| $\hat{q}$ (move probability) | 0.009***<br>(0.002) | 0.010***<br>(0.002) | 0.009***<br>(0.002) |
| $\hat{p}$ (error probability) | 0.153***<br>(0.006) | 0.162***<br>(0.005) | 0.157***<br>(0.008) |
| N (sibling pairs) | 6,507 | 8,343 | 3,632 |

Note: The sample is restricted to the first sibling pair observed in each family. Column (1) restricts the sample to female sibling pairs, (2) to mixed-gender sibling pairs, (3) to male sibling pairs. Heteroskedasticity robust standard errors are shown in parentheses. Standard errors for $\hat{p}$ were computed using the delta method. Significance levels are indicated as follows: \* p<0.1, \*\* p<0.05, \*\*\* p<0.01

**Table C.3:** Differences in siblings’ birth district and their age gap - across different district types.

|  | Different birth district: |  |  |
| --- | --- | --- | --- |
|  | (1)<br>Rural | (2)<br>Urban / municipal | (3)<br>Metropolitan |
| Age gap (years) | 0.021***<br>(0.004) | 0.008***<br>(0.001) | 0.009<br>(0.006) |
| Constant | 0.485***<br>(0.021) | 0.257***<br>(0.007) | 0.632***<br>(0.035) |
| **Derived probabilities:** |  |  |  |
| $\hat{q}$ (move probability) | 0.021***<br>(0.004) | 0.008***<br>(0.001) | 0.009<br>(0.006) |
| $\hat{p}$ (error probability) | 0.282***<br>(0.014) | 0.138***<br>(0.004) | 0.394***<br>(0.029) |
| N (sibling pairs) | 1,729 | 14,618 | 563 |

Note: The sample is restricted to the first sibling pair observed in each family. Each column of the table restricts the sample to sibling pairs in which the first (i.e. older) sibling is born in a district of the given type. Heteroskedasticity robust standard errors are shown in parentheses. Standard errors for $\hat{p}$ were computed using the delta method. Significance levels are indicated as follows: \* $p < 0.1$ , \*\* $p < 0.05$ , \*\*\* $p < 0.01$

**Table C.4:** Differences in siblings’ birth district - by district type.

| 1st sibling: | 2nd sibling: |  |  |  |  |
| --- | --- | --- | --- | --- | --- |
|  | Same district | Different district |  |  | Total |
|  |  | Rural | Urban / municipal | Metropolitan |  |
| Rural | 42.42%<br>(730) | 8.60%<br>(148) | 48.11%<br>(828) | 0.87%<br>(15) | 100.00%<br>(1,721) |
| Urban / municipal | 70.99%<br>(10,340) | 6.14%<br>(895) | 22.03%<br>(3,209) | 0.83%<br>(121) | 100.00%<br>(14,565) |
| Metropolitan | 33.03%<br>(184) | 4.67%<br>(26) | 35.19%<br>(196) | 27.11%<br>(151) | 100.00%<br>(557) |
| Total | 66.82%<br>(11,254) | 6.35%<br>(1,069) | 25.13%<br>(4,233) | 1.70%<br>(287) | 100.00%<br>(16,843) |

Note: The table shows row percentages and the corresponding frequency counts in parentheses.

**Table C.5:** Differences in siblings’ birth district and their age gap - across quartiles of the district population density.

|  | Quartiles of district-level population density: |  |  |  |
| --- | --- | --- | --- | --- |
|  | (1)<br>Q1 | (2)<br>Q2 | (3)<br>Q3 | (4)<br>Q4 |
| Age gap (years) | 0.014***<br>(0.002) | 0.012***<br>(0.002) | 0.006***<br>(0.002) | 0.006***<br>(0.002) |
| Constant | 0.376***<br>(0.013) | 0.358***<br>(0.012) | 0.233***<br>(0.012) | 0.211***<br>(0.010) |
| **Derived probabilities:** |  |  |  |  |
| $\hat{q}$ (move probability) | 0.014***<br>(0.002) | 0.012***<br>(0.002) | 0.006***<br>(0.002) | 0.006***<br>(0.002) |
| $\hat{p}$ (error probability) | 0.210***<br>(0.008) | 0.199***<br>(0.008) | 0.124***<br>(0.007) | 0.112***<br>(0.005) |
| N (sibling pairs) | 4,257 | 4,637 | 3,886 | 5,702 |

Note: The sample is restricted to the first sibling pair observed in each family. Each column of the table restricts the sample to sibling pairs in which the first (i.e. older) sibling is born in a district from the given population density quartile. Q1 denotes the 25% of districts with the lowest population density, Q4 the 25% of districts with the highest population density. Heteroskedasticity robust standard errors are shown in parentheses. Standard errors for $\hat{p}$ were computed using the delta method. Significance levels are indicated as follows: \* p<0.1, \*\* p<0.05, \*\*\* p<0.01

**Table C.6:** Differences in siblings’ birth district and their age gap - across quartiles of the district share of early school leavers.

|  | Quartiles of share left education by age 14: |  |  |  |
| --- | --- | --- | --- | --- |
|  | (1)<br>Q1 | (2)<br>Q2 | (3)<br>Q3 | (4)<br>Q4 |
| Age gap (years) | 0.014***<br>(0.002) | 0.009***<br>(0.002) | 0.010***<br>(0.002) | 0.007***<br>(0.002) |
| Constant | 0.365***<br>(0.012) | 0.264***<br>(0.013) | 0.228***<br>(0.012) | 0.280***<br>(0.011) |
| **Derived probabilities:** |  |  |  |  |
| $\hat{q}$ (move probability) | 0.014***<br>(0.002) | 0.009***<br>(0.002) | 0.010***<br>(0.002) | 0.007***<br>(0.002) |
| $\hat{p}$ (error probability) | 0.203***<br>(0.008) | 0.142***<br>(0.007) | 0.121***<br>(0.007) | 0.152***<br>(0.006) |
| N (sibling pairs) | 4,930 | 3,901 | 3,873 | 5,778 |

Note: The sample is restricted to the first sibling pair observed in each family. Each column of the table restricts the sample to sibling pairs in which the first (i.e. older) sibling is born in a district from the given quartile of the district-level share of those who left full-time education by age 14 according to the 1951 census<sup>25,26</sup>. Q1 denotes the 25% of districts with the lowest share of early school leavers, Q4 the 25% of districts with the highest share of early school leavers. Heteroskedasticity robust standard errors are shown in parentheses. Standard errors for $\hat{p}$ were computed using the delta method. Significance levels are indicated as follows: \* p<0.1, \*\* p<0.05, \*\*\* p<0.01

**Table C.7:** Differences in siblings’ birth district and their age gap - across quartiles of the polygenic index for education.

|  | Quartiles of PGI for education: |  |  |  |
| --- | --- | --- | --- | --- |
|  | (1)<br>Q1 | (2)<br>Q2 | (3)<br>Q3 | (4)<br>Q4 |
| Age gap (years) | 0.008***<br>(0.002) | 0.007***<br>(0.002) | 0.014***<br>(0.002) | 0.010***<br>(0.002) |
| Constant | 0.255***<br>(0.012) | 0.287***<br>(0.012) | 0.274***<br>(0.012) | 0.330***<br>(0.013) |
| **Derived probabilities:** |  |  |  |  |
| $\hat{q}$ (move probability) | 0.008***<br>(0.002) | 0.007***<br>(0.002) | 0.014***<br>(0.002) | 0.010***<br>(0.002) |
| $\hat{p}$ (error probability) | 0.137***<br>(0.007) | 0.156***<br>(0.007) | 0.148***<br>(0.007) | 0.181***<br>(0.008) |
| N (sibling pairs) | 4,512 | 4,512 | 4,512 | 4,512 |

Note: The sample is restricted to the first sibling pair observed in each family. Each column of the table restricts the sample to sibling pairs in which the mean PGI for education of the sibling pair lies in the given quartile of the distribution. Q1 denotes the 25% of sibling pairs with the lowest mean PGI, Q4 the 25% with the highest mean PGI. The PGI for education is from the PGI repository<sup>35</sup>. Heteroskedasticity robust standard errors are shown in parentheses. Standard errors for $\hat{p}$ were computed using the delta method. Significance levels are indicated as follows: \* p<0.1, \*\* p<0.05, \*\*\* p<0.01

**Table C.8:** Differences in siblings’ birth district and their age gap - by district shares of early school leavers and siblings’ polygenic index for education.

|  | Low education area |  | High education area |  |
| --- | --- | --- | --- | --- |
|  | (1)<br>Low PGI | (2)<br>High PGI | (3)<br>Low PGI | (4)<br>High PGI |
| Age gap (years) | 0.007***<br>(0.002) | 0.008***<br>(0.002) | 0.009***<br>(0.002) | 0.014***<br>(0.002) |
| Constant | 0.264***<br>(0.012) | 0.269***<br>(0.013) | 0.282***<br>(0.013) | 0.339***<br>(0.013) |
| **Derived probabilities:** |  |  |  |  |
| $\hat{q}$ (move probability) | 0.007***<br>(0.002) | 0.008***<br>(0.002) | 0.009***<br>(0.002) | 0.014***<br>(0.002) |
| $\hat{p}$ (error probability) | 0.142***<br>(0.007) | 0.145***<br>(0.008) | 0.153***<br>(0.007) | 0.187***<br>(0.008) |
| N (sibling pairs) | 4,290 | 3,710 | 3,984 | 4,562 |

Note: The sample is restricted to the first sibling pair observed in each family. Columns (1)-(2) of the table restrict the sample to sibling pairs in which the first (i.e. older) sibling is born in a district with an above-median share of those who left full-time education by age 14 according to the 1951 census<sup>25,26</sup>, columns (3)-(4) restrict the sample to sibling pairs in which the first sibling is born in a district with a below-median share of early school leavers. Columns (1) and (3) furthermore restrict the sample to sibling pairs in which the mean PGI for education of the sibling pair is below the median, columns (2) and (4) restrict to sibling pairs with an above-median PGI. The PGI for education is from the PGI repository<sup>35</sup>. Heteroskedasticity robust standard errors are shown in parentheses. Standard errors for $\hat{p}$ were computed using the delta method. Significance levels are indicated as follows: \* p<0.1, \*\* p<0.05, \*\*\* p<0.01

**Figure C.1:**
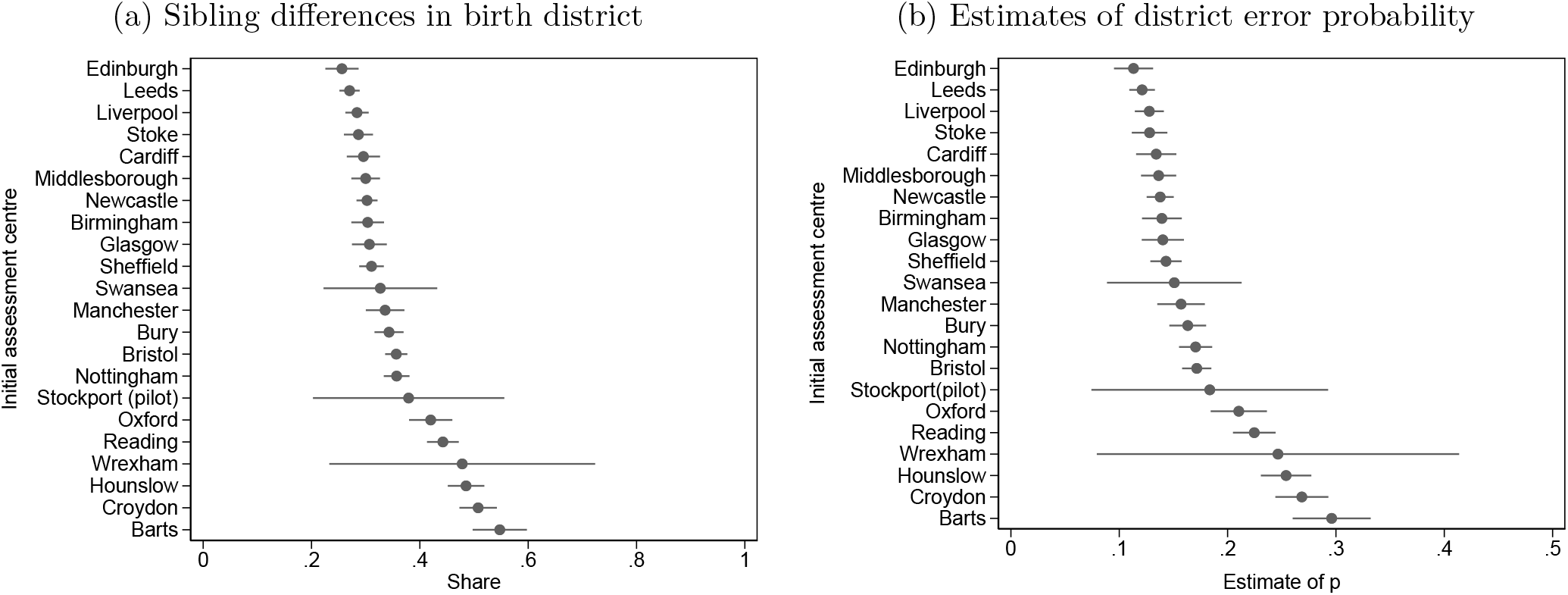
Accuracy of siblings’ birth district across assessment centre locations Note: This analysis was conducted at the individual level (using both siblings in a family), standard errors were clustered at the family level. In panel b), *q* was jointly estimated across all assessment centres. Bars represent 95% confidence intervals.

## Appendix D Attenuation bias - Additional results

**Figure D.1:**
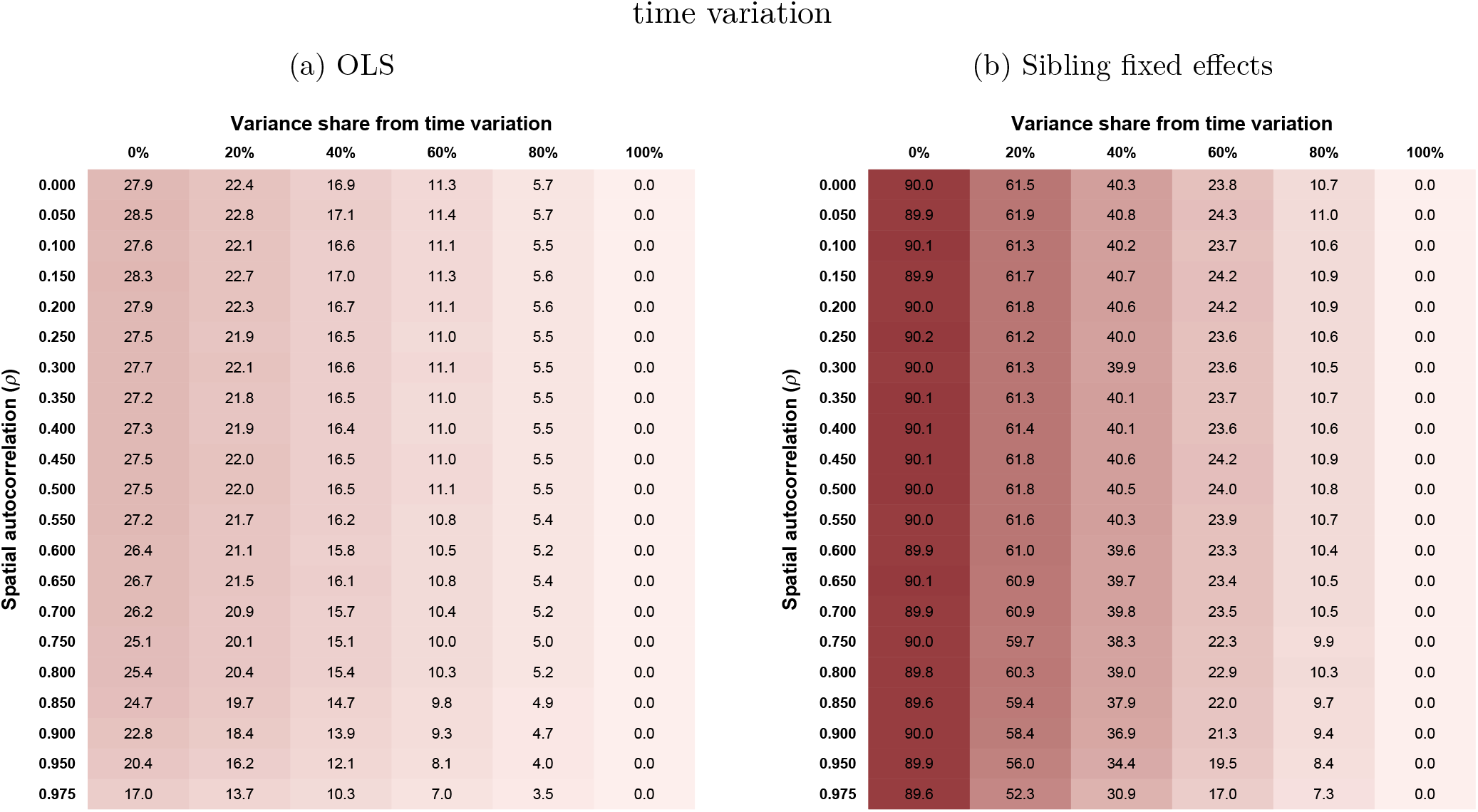
Attenuation bias for coordinate-level data with different levels of spatial correlation and time variation Note: The attenuation bias values shown are the mean bias (in %) from simulations of OLS and sibling fixed effects estimations with 1000 repetitions. For each level of spatial autocorrelation (*ρ*), 10 coordinate-level variables were simulated at a 1km resolution and merged to the sibling sample. The coordinate-level spatial variables were combined with normally distributed year-month of birth fixed effects to simulate time-varying spatial exposures. The columns of the tables correspond to different ratios of spatial to temporal variation when simulating the exposure variable - as indicated by the share of the exposure variance due to time variation. Each simulated variable was then used in 100 simulations of the attenuation bias based on an error probability for the location of birth *p* = 0.284 and a move probability *q* = 0.012.

**Figure D.2:**
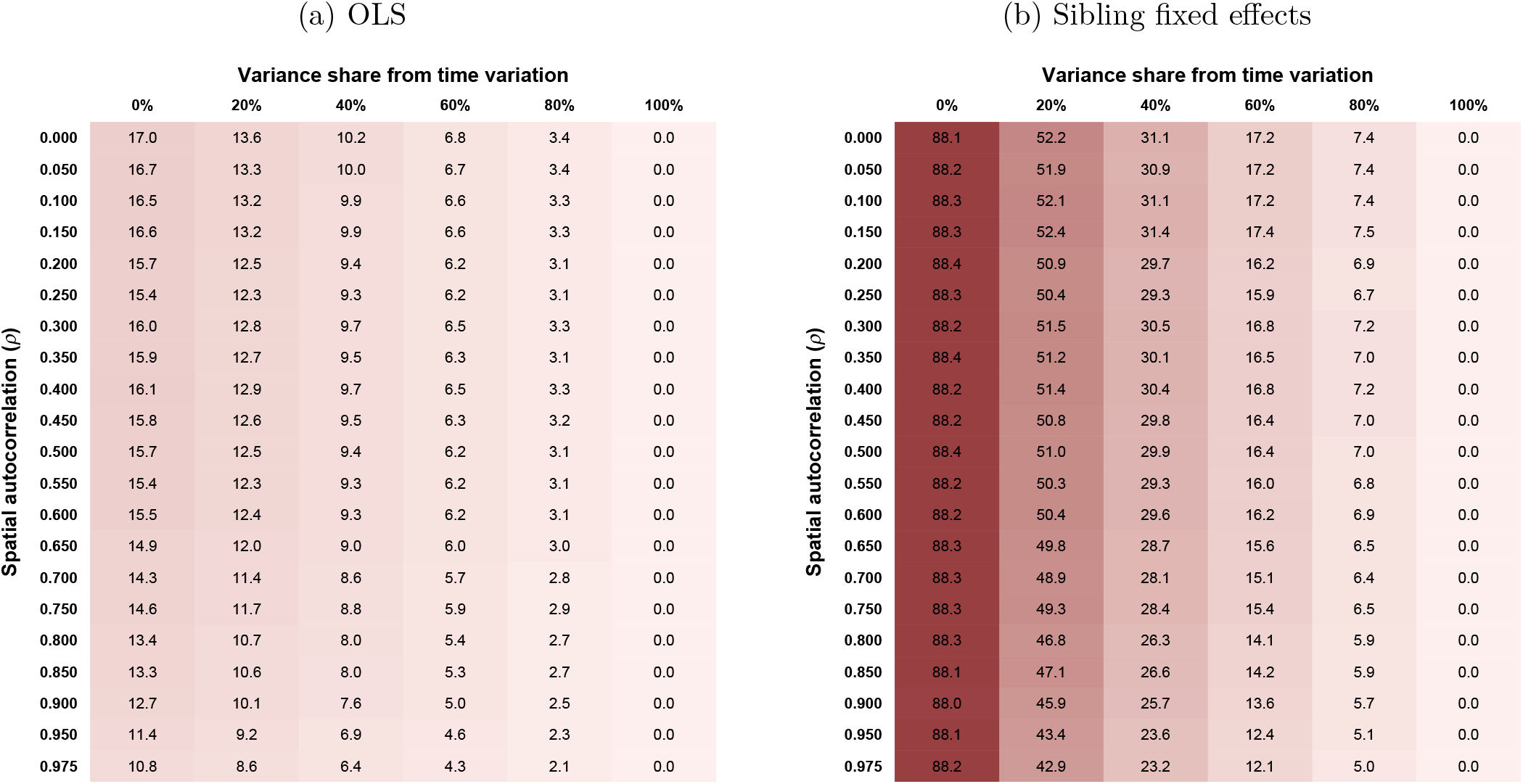
Attenuation bias for parish-level data with different levels of spatial correlation and time variation Note: The attenuation bias values shown are the mean bias (in %) from simulations of OLS and sibling fixed effects estimations with 1000 repetitions. For each level of spatial autocorrelation (*ρ*), 10 parish-level variables were simulated and merged to the sibling sample. The parish-level spatial variables were combined with normally distributed year-month of birth fixed effects to simulate time-varying spatial exposures. The columns of the tables correspond to different ratios of spatial to temporal variation when simulating the exposure variable - as indicated by the share of the exposure variance due to time variation. Each simulated variable was then used in 100 simulations of the attenuation bias based on an error probability for the parish of birth *p* = 0.168 and a move probability *q* = 0.009.

## Appendix E Spatial distribution of genetic principal components

**Figure E.1:**
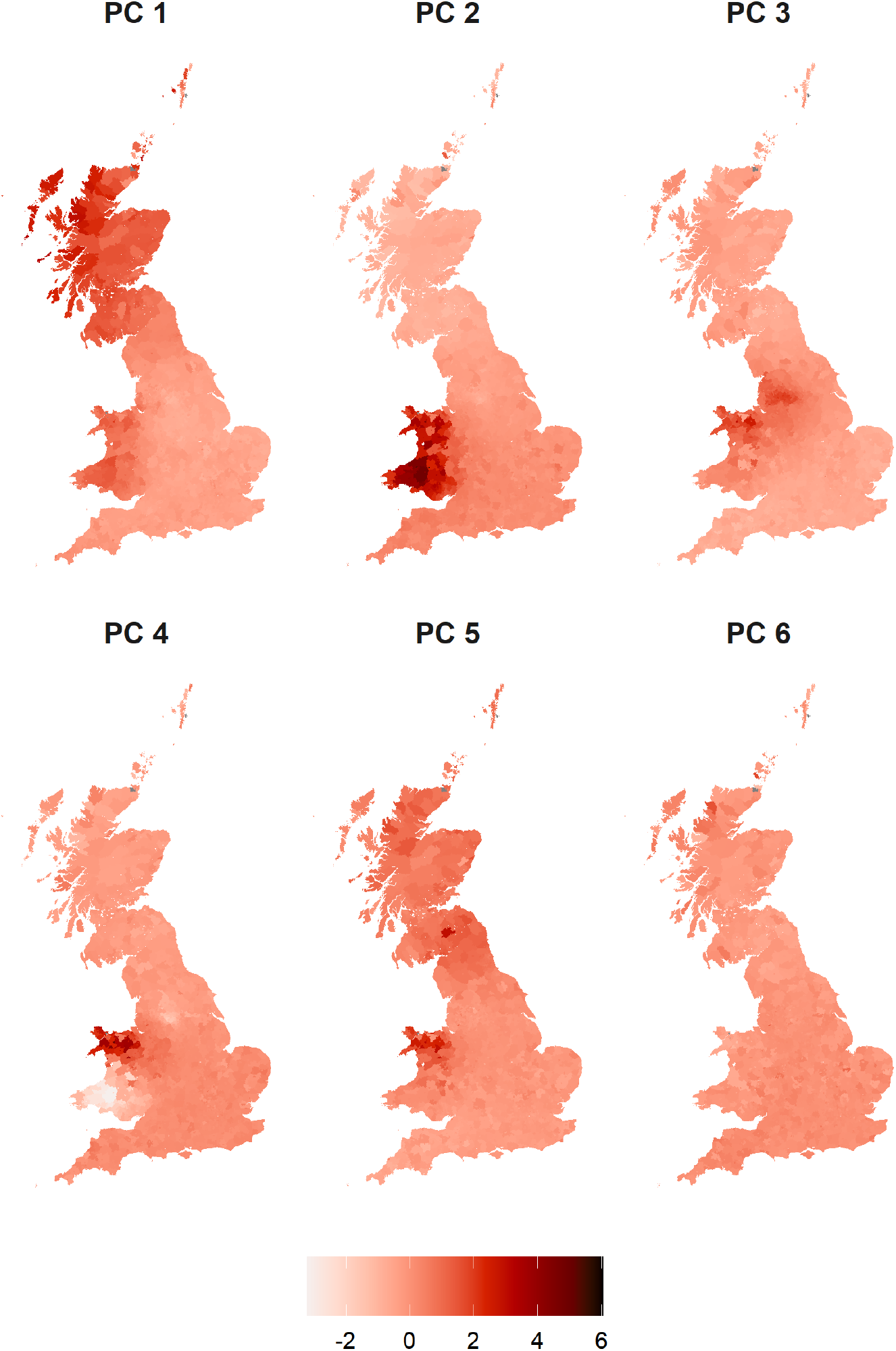
Geographical distribution of genetic principal components Note: The figures shows for each district the mean genetic principal components of UKB participants who reported being born in this district.

